# MYC-associated factor MAX is an essential regulator of the clock core network

**DOI:** 10.1101/771329

**Authors:** Olga Blaževitš, Nityanand Bolshette, Donatella Vecchio, Ana Guijarro, Ottavio Croci, Stefano Campaner, Benedetto Grimaldi

## Abstract

The circadian transcriptional network is based on a competition between transcriptional activator and repressor complexes regulating the rhythmic expression of clock-controlled genes. We show here that the MYC-Associated factor X, MAX, plays a repressive role in this network and operates through its MYC-independent binding to E-box-containing regulatory regions within the promoters of circadian BMAL1 targets. This clock function of MAX is essential for maintaining a proper circadian rhythm but separated by the role of MAX as a partner of MYC in controlling cell proliferation. We also identified MAX Network Transcriptional repressor, MNT, as a fundamental partner of MAX-mediated circadian regulation. Collectively, our data indicate that MAX is an integral part of the core molecular clock and keeps the balance between positive and negative elements of the molecular clock machinery. Accordingly, alteration of MAX transcriptional complexes may contribute to circadian dysfunction in pathological contexts.

## Introduction

Many cellular processes obey an endogenous cell-autonomous clock (the circadian clock) that has an intrinsic period of approximately 24 hours (Dibner et al., 2010; Reinke and Asher, 2019; Schibler and Sassone-Corsi, 2002). The molecular mechanism underlying these circadian rhythms is based on the interconnected transcriptional–translational feedback loops where specific transcription factors repress the expression of their own target genes (Ercolani et al., 2015; Ko and Takahashi, 2006; Takahashi, 2017).

Studies in cultured cells clearly showed the cell-autonomous feature of the transcriptional circadian rhythmicity and allowed the dissection of the molecular architecture of the clock (Lananna et al., 2018; Nagoshi et al., 2004).

Accordingly, the clock core network has been conceptualized as two transcriptional complexes operating in an antagonistic manner on the expression of clock-controlled genes (CCGs). On the one hand, the proteins CLOCK and BMAL1 interact to form a clock activator complex that stimulates the transcription of CCGs by recognizing E-box and E-box-like cis-regulative elements proximal to their core promoters (Shearman et al., 2000). On the other hand, the association of PERIOD and CRYPTOCHROME proteins with CLOCK/BMAL1 forms a transcriptional repressor complex that decreases CLOCK/BMAL1-dependent transcription (Cho et al., 2012; Van Der Horst et al., 1999; Vitaterna et al., 1999; Zheng et al., 2001). The periodic competition between clock-activator and clock-repressor complexes determines the circadian expression of around 5-10% of the mammalian transcriptome (Buhr and Takahashi, 2013; Miller et al., 2007; Panda et al., 2002). Nevertheless, additional negative regulators, such as REV-ERB nuclear receptors (Cho et al., 2012; Preitner et al., 2002), appears important for a proper circadian rhythm and mathematical modelling of the circadian clock gene-regulatory network indicated that a synergy of multiple inhibitions are required for robust self-sustained oscillations (Pett et al., 2016). In addition, a proper balance between activators and repressors of E-boxes has been proposed as a crucial requirement for generating circadian rhythms (Kim and Forger, 2012; Lee et al., 2011).

Disruption of the molecular clock is associated with a variety of human pathologies, including cancer (Ercolani et al., 2015; Roenneberg and Merrow, 2016). Ectopic overexpression of the oncogenic MYC protein has been recently reported to alter circadian gene expression in cancer cell lines, although the molecular mechanism behind this MYC function is still highly debated (Altman et al., 2017; Altman et al., 2015; Shostak et al., 2017; Shostak et al., 2016).

MYC is a transcription factor that can either activate or repress transcription depending on the interacting protein partners (Alderton, 2014; Carroll et al., 2018). As a heterodimer with the MYC-associated X-factor (MAX) protein, MYC stimulates transcription of diverse genes bearing promoter-proximal E-boxes, including important cell cycle and metabolic genes (Bretones et al., 2015; Wahlström and Henriksson, 2015) (Amati et al., 1992; Kretzner et al., 1992; Seoane et al., 2002). However, MYC can also repress gene expression when recruited in complex with MIZ1 to non-E-box sites in the promoters of MIZ1 target genes (Gebhardt et al., 2006; Peukert et al., 1997; Wiese et al., 2013).

Whether MYC-mediated alteration of the circadian rhythm depended on one or both mechanisms is still debated. Indeed, MYC overexpression was shown either to interfere with E-box driven transcription of the BMAL1-containing molecular clock complex (Altman et al., 2015) or to act as a direct transcriptional repressor of BMAL1 in an E-box-independent fashion (Shostak et al., 2016).

We report here that MAX operates as an unexpected integral and essential regulator of the circadian transcriptional network in a MYC-independent manner in both cancer and non-cancerous cell lines. We further identified the MAX binding protein, MNT, as a fundamental component of MAX-mediated clock regulation. Our data also implies that circadian disruption upon ectopic MYC overexpression would depend on perturbation of a physiological repression operated by MAX/MNT complex.

## Results

### Knockdown of MAX represses the transcription of core clock genes in cancer cell lines

Studies in U2OS cells over-expressing an ectopic MYC protein have shown that elevated levels of MYC can profoundly alter the expression of CLOCK/BMAL1-regulated genes (Altman et al., 2015; Shostak et al., 2016). Since up-regulation of MYC has been reported in many triple negative breast cancer (TNBC) cells (Fallah et al., 2017), we decided to evaluate whether this oncogene might control clock gene transcription in a TNBC cell line, MDA-MB-231. Thus, we compared mRNA levels of core clock genes in cells in which expression of *BMAL1*, *MYC* or *MAX* were knocked down by siRNA. In line with the dual role of the BMAL1-containing circadian complex in transcriptional regulation and with observations in *Bmal1*^−/−^ mice (Kondratov et al., 2006) (Hatanaka et al., 2010), the knock-down of BMAL1 reduced the levels of some clock transcripts (*PER1*, *NR1D1*, *NR1D2* and *TEF*) while de-repressed *PER2*, *CRY1* and *CRY2* transcription (Fig. 1A). Conversely, the expression of characterized MYC target genes involved in cell proliferation (Bretones et al., 2015), such as *CDK4*, *CDC25C*, *RCF4*, *NCL* and *MCM2*, showed negligible differences in *BMAL1*-silenced cells compared with control cells (Fig. 1A).

**Figure 1.**
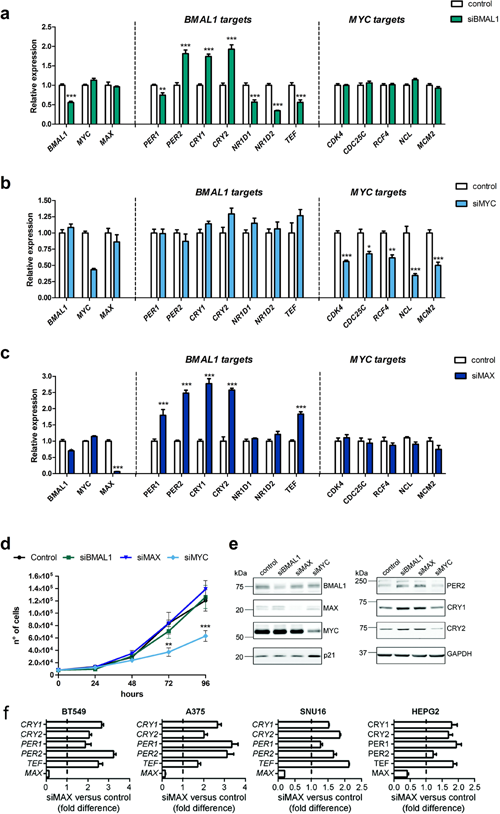
Knockdown of MAX alters the expression of core clock genes. (*A-C*) Expression of the circadian BMAL1 targets and the cell cycle MYC targets in MDA-MB-231 with knocked down *BMAL1* (siBMAL1), *MYC* (siMYC) or *MAX* (siMAX). A non-coding siRNA was used as control. Relative expression was determined by qRT-PCR using *GAPDH* for normalization. Values of control cells were set to 1. Shown as mean ± SEM, n ≥ 6. *P < 0.05. **P < 0.01 and ***P < 0.001, two-way ANOVA with Bonferroni post hoc test, silencing versus control. (*D*) Growth curve of MDA-MB-231 transfected with siRNA sequences against *BMAL1*, *MYC*, *MAX* or a non-targeting control. Shown as mean ± SEM, n ≥ 6. **P < 0.01 and ***P < 0.001, two-way ANOVA with Bonferroni post hoc test, siMYC versus control. (*E*) Immunoblot of protein samples from siBMAL1, siMYC, siMAX and control MDA-MB-231 cells with specific antibodies against the indicated proteins. GAPDH was used as a loading control. (*F*) The expression of *MAX*, *CRY1*, *CRY2*, *PER1*, *PER2* and *TEF* was analyzed in breast (BT549), skin (A375), stomach (SNU16) and liver (HEPG2) cancer cell lines knocked down for MAX. Values of control cells were set to 1 (dotted line). Shown as mean ± SEM, n = 3. See also related Supplemental Figure S1.

While knockdown of MYC markedly reduced mRNA levels of MYC proliferative-related targets, it had negligible effects on BMAL1-regulated genes (Fig. 1B). Strikingly, cells with knocked down MAX showed drastic alterations in diverse clock transcripts (Fig. 1C). Indeed, the majority of core clock genes were significantly up-regulated upon MAX silencing, including the repressor genes, *PER1*, *PER2*, *CRY1* and *CRY2* (Fig. 1C). Notably, the decline of MAX in *MAX*-silenced cells was not sufficient for significantly influencing proliferative MYC targets, suggesting that the remaining MAX protein could still ensure a proper function of the MAX/MYC complex.

In line with our transcriptional data, knockdown of either *MAX* or *BMAL1* had no effect on cell proliferation, while MYC-silenced MDA-MB-231 cells showed significantly reduced growth compared with control cells (Fig. 1D). Immunoblot analysis in MDA-MB-231-silenced cells confirmed that reduction of either BMAL1 or MAX elevated PER2, CRY1 and CRY2 protein levels, whereas knockdown of MYC had no such effect (Fig. 1E). Consistent with the observation that only MYC silencing influenced MDA-MB-231 proliferation, protein levels of the cell cycle regulator Cyclin Dependent Kinase Inhibitor 1A (CDKN1A, also known as p21) showed differences solely in MYC-silenced cells (Fig. 1E).

Knockdown of *MAX* expression by using two additional diverse and non-redundant siRNA sequences against *MAX* transcripts similarly affected *BMAL1*, *PER1*, *PER2*, *CRY1*, *CRY2*, and *TEF* transcript levels (Fig. S1), thus ruling out that altered clock gene expression in MAX-silenced cells derived from off-target effects.

Notably, the knockdown of MAX significantly increased the expression of clock genes in a different TNBC cell line, BT549, as well as in cancer cell lines originated from skin (A375), stomach (SNU16) and liver (HEPG2) tumors (Fig. 1F), thus indicating that MAX-mediated regulation of clock genes might be extended to diverse human cell lines.

### MAX-inhibition of the core clock genes is independent from CRY-mediated repression, but requires a functional E-box responsive element

The effect of MAX silencing on the transcription of clock genes resembles the molecular phenotype observed in cells or tissues lacking CRYs repressor proteins (i.e. transcriptional de-repression of CLOCK/BMAL1/CRYs targets) (Kondratov et al., 2006; Takahashi, 2017). We thus investigated whether MAX might affect CRY-mediated repression by evaluating the expression of *PER1* and *PER2* following treatment with a CRY agonist (KL001 (Hirota et al., 2012)) in MDA-MB-231 knocked down for BMAL1 or MAX. As expected, KL001 augmented CRY transcriptional repression in control cells, as indicated by the significant decrease of *PER1* and *PER2* mRNA levels in KL001 treated cells compared with vehicle (Fig. 2A and B). In line with the essential role of CLOCK/BMAL1 complex in mediating CRY1 activity (Takahashi, 2017), the knockdown of *BMAL1* strongly reduced KL001-mediated inhibition of *PER1* and *PER2* transcription.

**Figure 2.**
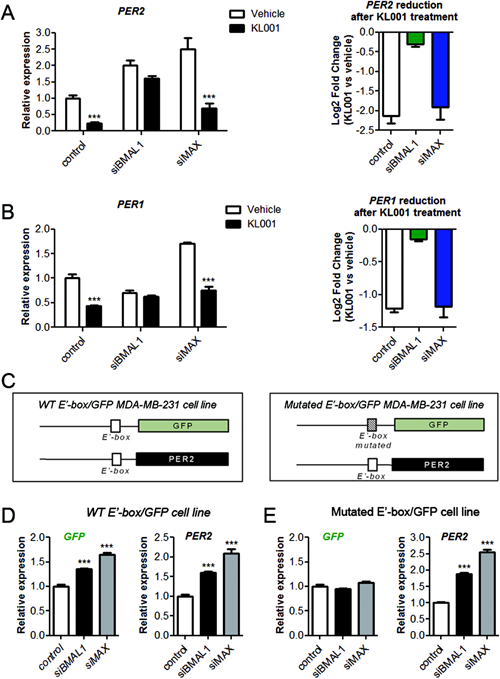
MAX-inhibition of the core clock genes is independent from CRY-mediated repression, but requires a functional E-box responsive element.(*A and B*) MDA-MB-231 cells transfected with siRNA sequences against *BMAL1*, *MAX* or a non-targeting control were treated 24 h with vehicle (DMSO) or 10 μM CRY1 agonist (KL001). The effect of KL001 on the expression of CRY1 targets, *PER1* and *PER2*, was evaluated by qRT-PCR using *GAPDH* for normalization. Values of untreated control cells were set to 1. Shown as mean ± SEM, n = 3. ***P < 0.001, two-way ANOVA with Bonferroni post hoc test, vehicle versus KL001. Right panel reports the log2 ratio between KL001 and vehicle samples, thus showing reduction of *PER1* or *PER2* expression after the treatment with CRY agonist. (*C*) Schematic representation of two MDA-MB-231 reporter cell lines bearing the sequence coding for the Green Fluorescent Protein (GFP) gene controlled by a promoter fragment of *PER2* containing a wild-type or a mutated version of the clock regulated E’-box element (*WT E’-box/GFP* and *Mutated E’-box/GFP* cells, respectively). (*D and E*) Expression of the endogenous *PER2* gene and the GFP driven by a *WT E’-box* or a *mutated E’-box* in reporter cells with knocked down BMAL1 (siBMAL1) or MAX (siMAX). A non-coding siRNA was used as control. Relative expression was determined by qRT-PCR using *GAPDH* for normalization. Values of control cells were set to 1. Shown as mean ± SEM, n = 4. ***P < 0.001, two-way ANOVA with Bonferroni post hoc test, silencing versus control. See also related Supplemental Figure S2.

In contrast, KL001 efficacy was preserved upon *MAX* knocked down, as indicated by a comparable drug-related decrease in *PER* transcripts in both MAX-silenced and control cells (Fig. 2A and B). In addition, MAX silencing significantly enhanced the transcription of PER1, PER2, CRY2, and TEF in both control and CRY1-silenced cells (Fig. S2A).

Collectively, our results suggest that MAX repression operates independently from the activity of the BMAL1/CLOCK/CRY repressor complex.

The above results do not preclude the possibility that BMAL1 and MAX might regulate the expression of clock target genes by acting on similar regulatory regions. Indeed, both these transcription factors interact with E-box and E-box-like elements (Hardin, 2004; Lüscher, 2001). To evaluate this aspect, we generated two transgenic MDA-MB-231 cell lines expressing the Green Fluorescent Protein (*GFP*) gene under the control of a promoter fragment of *PER2* containing either a wild-type or a mutated E’-box element (*E’-box-GFP and E’*^*mut*^*-box-GFP* cells, respectively) (Fig. 2C). Indicating a functional clock regulation of our cell-based reporter system, KL001 reduced the expression of *GFP* in *E’-box-GFP*, but not in *E’*^*mut*^*-box-GFP* cells (Fig. S2B). As an internal control, KL001 treatment inhibited the transcription of the endogenous *PER2* gene in both cell lines. We thus evaluated the effect of BMAL1 or MAX silencing in the GFP reporter cells. Revealing that both factors require a functional E’-box for their transcriptional activity, the knockdown of either BMAL1 or MAX reduced GFP expression in *E’-box-GFP*, but not in *E’-box*^*Mut*^*-GFP* cells (Fig. 2D and E). In contrast, endogenous *PER2* transcription was elevated in both cell lines upon silencing of either BMAL1 or MAX.

### MAX is recruited on BMAL1 bound genomic regions in a MYC-independent manner

Our results with the GFP-reporter cell lines suggest that BMAL1 and MAX might be recruited on the same E-box containing regulatory regions within the promoters of clock target genes. To explore this possibility, we performed Chromatin Immuno-Precipitation sequencing (ChIP-seq) experiments with specific antibodies against BMAL1 and MAX proteins. This analysis revealed a large number of genomic regions bound by MAX (around 13000 peaks), while BMAL1 binding was limited to about 800 regions (Fig. 3A, Table S1 and S2). BMAL1 and MAX bound regions comprised both promoters and distal sites. Coherently with the circadian role of BMAL1 and its preference for E-box-containing sites, ontological annotation of BMAL1 bound regions showed a significant enrichment for circadian regulated genes and E-box motifs (Fig. S3A). Remarkably, 85% of the BMAL1 bound sites overlapped with MAX bound regions (Fig. 3B, C). Furthermore, the enrichment of MAX was significantly higher on the genomic regions bound by BMAL1 than on the loci sites in which BMAL1 was not present (Fig. 3D), thus indicating that BMAL1 target sites are bound by MAX with high affinity.

**Figure 3.**
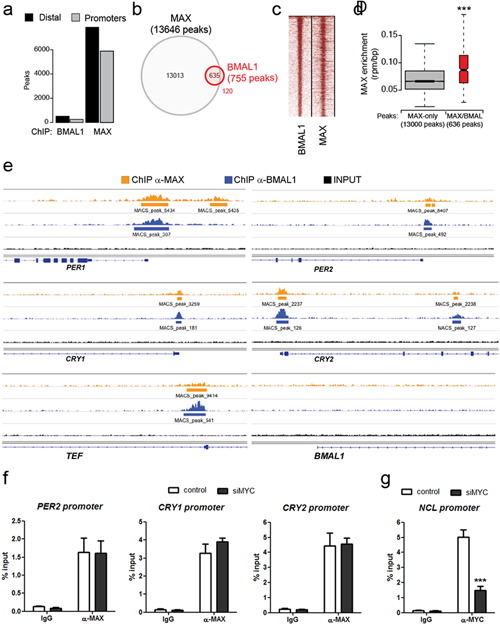
MAX is recruited on BMAL1-bound genomic regions in a MYC-independent manner. (A) Number of peaks identified for ChIP-seq of BMAL1 and MAX in MDA-MB-231 cells. Sub-setting of distal or promoter peaks was based on their proximity to annotated transcriptional starting site. (B) Venn diagram showing the overlap of ChIP-seq peaks of BMAL1 and MAX. (*C*) Heatmap of ChIP-seq signals of genomic regions co-bound by BMAL1 and MAX. (*D*) Box-plot showing the enrichment of MAX in genomic regions bound by MAX (MAX only) or by both MAX and BMAL1 (MAX/BMAL1). ***P<0.001, two tailed student t test. (*E*) Genomic snapshots of the promoter region of core circadian clock genes (*PER1*, *PER2*, *CRY1*, *CRY2*, *TEF*, *BMAL1*) showing the enrichment of MAX (orange) and BMAL1 (blue). (*F*) Recruitment of MAX on the E-box-containing promoters of *PER2*, *CRY1* and *CRY2* in MYC-silenced and control MDA-MB-231 cells. Enrichment of MAX was evaluated by quantitative PCR of immunoprecipitated DNA compared with input DNA (% of input). Immunoglobulin G (IgG) was used as a negative control. Shown as mean + SEM, n =3. (*G*) Chromatin samples from (F) were immunoprecipitated with α-MYC antibody to confirm the actual reduction of MYC recruitment on a MYC-target gene (*NCL*) in MYC-silenced cells. ***P<0.001, two tailed student t test. See also related Supplemental Figure S3 and Table S1-S2.

Strikingly, MAX was present on the promoters of all the E-box-containing circadian factors up regulated upon MAX silencing (*PER1*, *PER2*, *CRY1*, *CRY2*, and *TEF*) (Fig. 3E), supporting a direct transcriptional repressive activity of MAX on the clock molecular machinery.

While MAX was not detected on the BMAL1 locus (Fig. 3E), we observed an enrichment of MAX on the promoters of NR1D1 and NR1D2 (Fig. S3B), which form a well-established feedback loop with BMAL1 in the circadian signalling network (Takahashi, 2017). It is thus conceivable that transcriptional de-repression of NR1D1/NR1D2 in MAX-silenced MDA-MB-231 was not observed because of the resulting compensatory lowering in BMAL1 levels (Fig. 1C). Consistent with this hypothesis, the knockdown of MAX in BMAL1-silenced cells significantly increased both *NR1D1* and *NR1D2* transcription (Fig. S3C). We further evaluated whether the recruitment of MAX on BMAL1 target promoters might be independent from MYC by immunoprecipitation of MYC-silenced and control chromatin samples with an anti-MAX antibody. Strikingly, MYC silencing resulted in negligible differences in the enrichment of MAX on the promoters of *PER2*, *CRY1* and *CRY2* (Fig. 3F). Confirming the actual reduction of MYC in MYC-silenced cells, ChIP with an anti-MYC antibody showed a drastic reduction of MYC recruitment on the *NCL* promoter in cells knocked down for MYC, compared with control (Fig. 3G).

Altogether, our data reveal that MAX can operate as a direct repressor of core clock genes in a MYC-independent manner.

### MAX and BMAL1 regulates the expression of common transcripts

Genome wide co-occurrence of BMAL1 and MAX on transcriptional regulatory regions suggest that these proteins might control the expression of common targets. To address this, we used a next generation sequencing (NGS) approach for the identification of differentially expressed genes (DEGs) in MAX- or BMAL1-silenced MDA-MB-231 cells. Transcript assembly and quantification of RNA-sequencing reads identified 4863 and 4247 differentially expressed genes (DEGs) upon knockdown of either BMAL1 or MAX, respectively (Supplemental Table 3-4). The comparison of the two sets of genes revealed that 2391 of siBMAL1 DEGs (almost 50%) were also differentially expressed in MAX-silenced cells (Fig. 4A and Supplemental Table 5). Within this subset, 662 transcripts showed a logarithmic fold change (LogFC) greater than 0.5. We thus analysed their co-occurrence in KEGG pathways for evaluating the transcriptional signalling more affected by both MAX and BMAL1 using a false discovery rate (FDR) q value < 0.01 as a cut-off. Strikingly, this analysis identified the circadian rhythm as a highly significant pathway (q<0.00001), together with focal adhesion and glycosaminoglycan degradation (Fig. 4B). Consistent with our quantitative RT-PCR experiments, the list of transcripts co-regulated by MAX and BMAL1 included *period* and *cryptochrome* circadian repressor genes (Supplemental Table 5). In addition, another negative regulator of BMAL1/CLOCK-mediated transcription, *BHLHE41* (also known as *DEC2*) (Honma et al., 2002), resulted under the control of both MAX and BMAL1.

**Figure 4.**
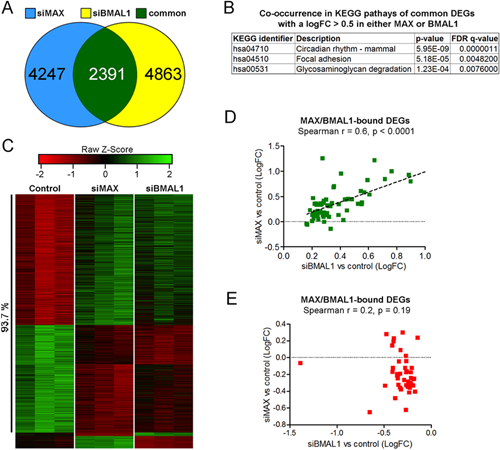
MAX and BMAL1 regulates the expression of common transcripts. (*A*) Venn diagram for differentially expressed genes (DEGs) in MDA-MB-231 cells with knocked down MAX (siMAX) or BMAL1 (siBMAL1). (*B*) Co-occurrence in KEGG pathways of the set of common siMAX and siBMAL1 DEGs with an absolute logFC > 0.5 upon the knockdown of either MAX or BMAL1. (*C*) Clustered heat map of triplicate normalized counts from common siMAX:siBMAL1 DEGs. The percentage of genes coherently altered in both MAX- and BMAL-silenced cells compared with control is shown. (*D*) Significant correlation between MAX/BMAL1-bound genes up-regulated in BMAL1-silenced and MAX-silenced cells. (*E)* Lack of correlation between MAX/BMAL1-bound genes down-regulated in BMAL1-silenced and MAX-silenced cells. See also related Supplemental Table S3-S5.

Notably, heat map and clustering analysis of DEGs in siMAX and siBMAL1 cells revealed that more than 90% of these genes (2241 out of 2391) were coherently altered in both conditions (i.e. their expression was altered in the same direction upon either MAX or BMAL1 silencing) (Fig. 4C). These results suggest that the overall effect of a reduction of MAX in silenced cells is a derepression of genes that are negatively regulated by the BMAL1-containing clock repressor complex. Supporting this hypothesis, LogFC values of BMAL1/MAX-bound genes upregulated by BMAL1-silencing positively correlated with their corresponding LogFC values in MAX-silenced cells (Fig. 4D). In contrast, no significant correlation was observed between BMAL1/MAX-bound genes downregulated upon the knockdown of BMAL1 (Fig. 4E).

Collectively, our data indicate that MAX is part of the negative arm of the molecular clock machinery.

### MAX is required for circadian gene expression

The above results strongly suggest that MAX might have a direct role in circadian transcriptional regulation, thus contributing to rhythmic oscillatory expression of clock target genes. To investigate this aspect, we silenced either BMAL1 or MAX in MDA-MB-231 cells expressing a firefly luciferase reporter controlled by the *BMAL1* promoter and monitored luciferase activity with a real-time luminometer after circadian synchronization by dexamethasone treatment (Ramanathan et al., 2012) (Fig. 5A). Baseline-subtracted luminescence data were then fitted to a sine wave and plotted to compare rhythmic patterns (Fig. 5B). This analysis showed a rhythmic oscillation of luminescence in dexamethasone-treated control cells over a 72 h period. Confirming that this rhythm was under the control of the circadian clock machinery, knockdown of *BMAL1* prevented the oscillation of the luciferase reporter. Strikingly, silencing of *MAX* markedly reduced rhythm amplitude compared with control cells (Fig. 5B).

**Figure 5.**
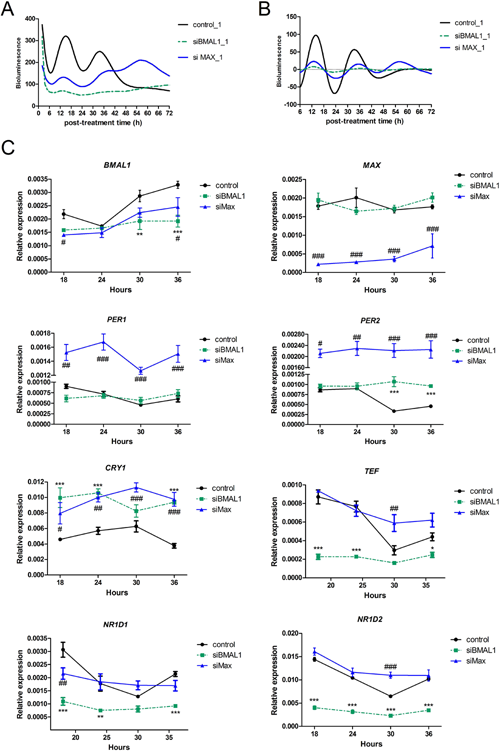
MAX is required for circadian gene expression in MDA-MB-231 cells. (*A*) Bioluminescence counts from circadian synchronized MDA-MB-231 cells expressing a firefly luciferase reporter controlled by the circadian-responsive *BMAL1* promoter with knocked down BMAL1 (siBMAL1) or MAX (siMAX). Cells transfected with a non-coding control was used as a control. (*B*) Baseline-subtracted luminescence data from (A) were fitted to a sine wave and plotted to compare rhythmic patterns. The first 6h upon synchronization were removed for the analysis to avoid interference of the dexamethasone response in the early time of synchronization. (*C*) Time-related expression of endogenous clock controlled genes in siBMAL1, siMAX and control synchronized MDA-MB-231 cells. Relative expression at the indicated dexamethasone post-treatment time points was determined by qRT-PCR using *GAPDH* for normalization. Shown as mean + SEM, n =3. *P<0.05, **P<0.01 and ***P<0.01, siBMAL1-silenced versus control cells. ^#^P<0.05, ^##^P<0.01 and ^###^P<0.001, siMAX-silenced versus control cells, two-way ANOVA with Bonferroni post hoc test.

We then analyzed the circadian profile of different endogenous clock genes in *BMAL1* and *MAX*-silenced cells synchronized by dexamethasone treatment. In line with observations in liver from *Bmal1*^−/−^ mice (Hatanaka et al., 2010; Kondratov et al., 2006), knockdown of *BMAL1* prevented time-dependent variations of all circadian transcripts (Fig. 5C).

Consistent with our luminescence analysis, *MAX*-silenced cells showed a reduced *BMAL1* oscillatory expression compared with control cells. Furthermore, reduction of *MAX* altered the expression of all circadian genes analyzed (Fig. 5C). Of note, *PER1*, *PER2* and *CRY1* transcript levels were constitutively higher at all time-points in *MAX*-silenced cells, which is consistent with their elevated expression in non-synchronous cells upon knockdown of *MAX*.

These results show that MAX is an essential regulator of circadian gene expression in MDA-MB-231 cells.

Cancer cell lines are associated with many mutations (Barretina et al., 2012), which might alter the activity and specificity of transcription factors. We thus evaluated whether MAX could control clock gene transcription in two non-cancerous human cell lines, foreskin fibroblast BJ-5ta and epithelial MCF10A. Similar to our observations in diverse cancer cells lines (Fig. 1F), MAX silencing increased the levels of clock transcripts in both BJ-5ta and MCF10A (Fig. 6A and B). Quantitative ChIP assays with an anti-MAX antibody further revealed that MAX-mediated clock gene expression in MCF10A also corresponded with a direct recruitment of MAX on the promoter of those circadian genes (Fig. 6C). Moreover, MAX silencing in *BMAL1-luc* MCF10A cells markedly reduced the amplitude of luciferase oscillation following dexamethasone synchronization treatment compared with control (Fig. 6D).

**Figure 6.**
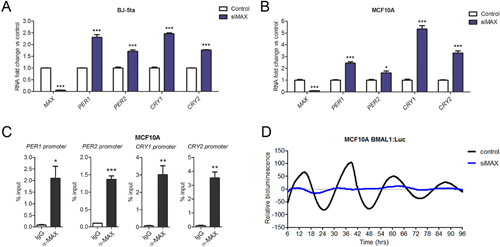
MAX-mediated circadian gene expression also operates in non-cancerous cells. (*A-B*) Expression of *MAX*, *PER1*, *PER2*, *CRY1*, and *CRY2* genes upon MAX silencing in foreskin fibroblast BJ-5ta and epithelial MCF10A cell lines. Relative expression was determined by qRT-PCR using *GAPDH* for normalization. Values of control cells were set to 1. Shown as mean + SEM, n ≥ 3. *P < 0.05. **P < 0.01 and ***P < 0.001, two-way ANOVA with Bonferroni post hoc test, silencing versus control. (*C*) Recruitment of MAX on the promoter of *PER1*, *PER2*, *CRY1* and *CRY2* in MCF10A cells. Enrichment of MAX was evaluated by quantitative PCR of immunoprecipitated DNA compared with input DNA (% of input). Immunoglobulin G (IgG) was used as a negative control. Shown as mean + SEM, n =3. *P < 0.05. **P < 0.01 and ***P < 0.001, unpaired two-tailed T-test. (*D*) Real-time bioluminescence oscillatory pattern in MAX-silenced and control MCF10A cells expressing a firefly luciferase reporter controlled by the circadian-responsive *BMAL1* promoter. Shown as baseline-subtracted luminescence data fitted to a sine wave.

Collectively, our data support an essential function of MAX in regulating the circadian clock of both cancer and non-cancerous cells.

### MAX dependent repression of clock genes requires MNT

MAX can operate as either an activator or a repressor of transcription depending on the interacting partner proteins (Kretzner et al., 1992; Nair and Burley, 2003). Among these protein complexes, those formed by MAX and MNT actively repress the expression of genes containing E-box elements in their regulatory regions (Terragni et al., 2011). Suggesting a role of MNT in MAX-mediated clock transcription, PER2 protein levels increased upon the knockdown of either MAX or MNT in MDA-MB-231 cells (Fig. 7A and B).

**Figure 7.**
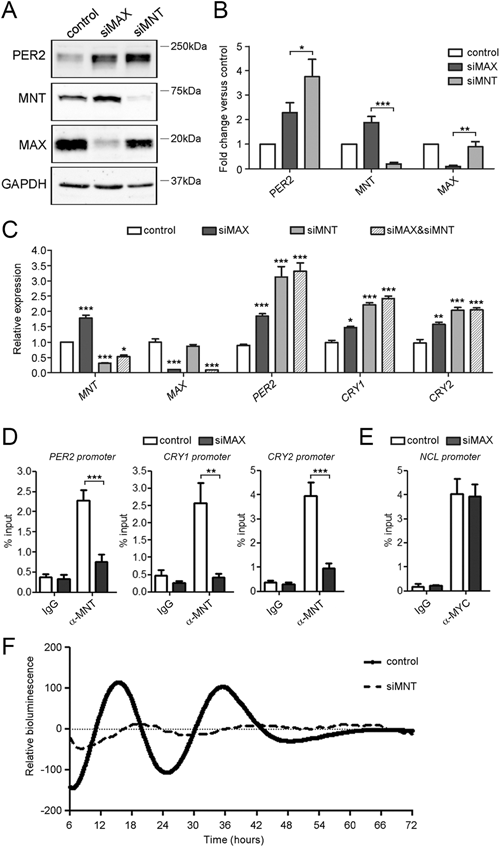
MAX dependent repression of clock genes requires MNT. (*A*) Protein levels of PER2, MNT and MAX in MDA-MB-231 cells transfected with siRNA sequences against MAX (siMAX), MNT (siMNT) or a non-targeting control. GAPDH was used as a loading control. (*B*) Densitometry analysis of protein signals for samples treated as in (A) is reported as relative protein levels normalized by GAPDH. Values of control cells were set to 1. Shown as mean + SEM, n=3. *P < 005, **P <0.01 and *P < 0.001, unpaired two-tailed T-test, siMAX versus siMNT. (*C*) Expression of clock genes *PER2*, *CRY1* and *CRY2* in MDA-MB-231 cells upon the knockdown of MAX (siMAX), MNT (siMNT), and both MAX and MNT (siMAX&siMNT). Relative expression was determined by qRT-PCR using *GAPDH* for normalization. Values of control cells were set to 1. Shown as mean + SEM, n ≥ 3. *P<0.05, **P<0.01 and ***P<0.01, two-way ANOVA with Bonferroni post hoc test, silenced versus control cells. (*D*) Recruitment of MNT on the promoters of PER2, CRY1 and CRY2 in MAX-silenced (siMAX) and control MDA-MB-231 cells. Enrichment of MNT was evaluated by quantitative PCR of immunoprecipitated DNA compared with input DNA (% of input). Immunoglobulin G (IgG) was used as a negative control. Shown as mean + SEM, n =3. ***P<0.001, two-way ANOVA with Bonferroni post hoc test, α-MNT precipitated DNA from siMAX versus control chromatin samples. (*E*) Chromatin samples from *D* (MAX-silenced and control) were immunoprecipitated with α-MYC. Enrichment of MYC on the MYC-target *NCL* promoter was evaluated by quantitative PCR of immunoprecipitated DNA compared with input DNA (% of input). Immunoglobulin G (IgG) was used as a negative control. Shown as mean + SEM, n =3. (*F*) Real-time bioluminescence oscillatory pattern in MNT-silenced (siMNT) and control MDA-MB-231 cells expressing a firefly luciferase reporter controlled by the circadian-responsive *BMAL1* promoter. Shown as baseline-subtracted luminescence data fitted to a sine wave. See also related Supplemental Figure S4.

We then evaluated the transcription of *PER2*, *CRY1*, and *CRY2* in MDA-MB-231 cells knocked down for *MAX*, *MNT* or both proteins. Consistent with our immunoblot analysis, MNT expression significantly increased in MAX-silenced cells (Fig. 7C). Strikingly, MAX reduction in siMAX/siMNT cells did not further alter the de-repression of clock genes observed with the single knockdown of MNT, indicating that a repressive complex formed by MAX and MNT regulates clock gene expression. Supporting this hypothesis, ChIP analysis in MAX-silenced MDA-MB-231 cells with an anti-MNT antibody revealed a MAX-mediated recruitment of MNT on *PER2*, *CRY1* and *CRY2* promoters (Fig. 7D). Notably, immunoprecipitation from the same MAX-silenced chromatin samples with an anti-MYC antibody revealed that the remaining MAX protein was still sufficient for a substantial binding of MYC on the promoter of NCL (Fig. 7E), which is consistent with the fact that MAX reduction did not affect MYC dependent transcription in MDA-MB-231 cells.

Similar to MAX silencing, knockdown of MNT in *BMAL1*-luc MDA-MB-231 cells markedly reduced luciferase oscillation after dexamethasone synchronization (Fig. 7F), further supporting a fundamental role of MNT in MAX-dependent clock regulation.

Indicating that this MNT function was not limited to breast cancer MDA-MB-231, MCF10A cells knocked down for MNT showed increased expression of *PER1*, *PER2*, *CRY1*, *CRY2*, and impaired oscillation of the luciferase circadian reporter (Fig. S4).

## Discussion

Our knockdown experiments indicate that MAX regulates the transcription of diverse genes belonging to the core clock machinery in both cancer and non-cancerous cell lines. Remarkably, knockdown of MAX was not sufficient to affect both the transcription of cell cycle-related MYC targets and MDA-MB-231 proliferation. Suggesting that the residual MAX protein still allowed for a MYC-dependent transcription of cell cycle genes, the recruitment of MYC on NCL promoter was not significantly reduced in MAX-silenced cells. These data imply that MAX-mediated activity on clock genes is independent from the role of MAX as a MYC-associated factor. Supporting this concept, the knockdown of MYC produced negligible effects on the expression of core clock genes and it did not alter the recruitment of MAX on *PER2*, *CRY1* and *CRY2* promoters.

MAX actively represses numerous core clock genes by its direct binding to E-box containing regions, as indicated by our ChIP-seq and qChIP analyses and by a lack of MAX-mediated derepression of a *PER2* promoter bearing a mutated E’-box sequence. Our genome-wide approaches further revealed that a reduction of MAX levels affects numerous BMAL1 regulated genes.

Notably, the core clock genes directly targeted and repressed by MAX include all the repressors belonging to both the primary (CRYs and PERs) and the accessory (NR1D1 and NR1D2) negative circadian feedback loops (Lee et al., 2011; Takahashi, 2017). Consequently, knockdown of MAX strongly altered the ratio between positive and negative elements of the molecular clock machinery in diverse cell lines. A proper stoichiometric balance between activators and repressors of E-boxes has been proposed as a crucial requirement for generating circadian rhythms (Kim and Forger, 2012; Lee et al., 2011). Accordingly, MAX silencing impaired rhythmic gene expression in both MDA-MB-231 and MCF10A cells, indicating an essential role of MAX in maintaining a functional circadian rhythm.

Collectively, our data support a model in which MAX is an essential part of the molecular clock and keeps the balance between clock activators and repressors.

We also identified MNT as a partner in MAX-mediated circadian regulation that is recruited on the promoters of clock core genes in a MAX-dependent manner. Our double knockdown experiments in MDA-MB-231 cells support a fundamental role of MNT in the repression of the molecular clock by MAX. In addition, MNT silencing strongly impaired circadian oscillation in both MDA-MB-231 and MCF10A cells. However, these results do not preclude the possibility that other heterodimerization partners of MAX (Hurlin and Huang, 2006) could contribute to the circadian clock in different cells or conditions (e.g. tissue development and differentiation), depending on the dynamics of MAX interactions (Carroll et al., 2018).

Our data also imply that circadian alteration upon MYC overexpression (Altman et al., 2015; Shostak et al., 2016) would depend on perturbation of a physiological repression operated by MAX/MNT complex. Indeed, it is well known that forced expression of MYC can antagonize with MNT for MAX binding to form a MAX/MYC activator complex (Carroll et al., 2018; Grandori et al., 2000). Consistent with this view, increased MYC levels in U2OS up-regulated clock genes which promoters showed a direct recruitment of MAX and MNT in our ChIP experiments (i.e. *PERs*, *CRYs* and *REV-ERBs*) (Altman et al., 2015). The presented data also indicate that alteration of MAX transcriptional network may contribute to circadian dysfunctions observed in several pathological contexts. For instance, downregulation of MNT has been recently proposed as a functional important event for the hypoxia response in a wide variety of injury and disease settings (Yang and Hurlin, 2017), and severe consequences caused by acute hypoxia have been correlated with defects in circadian rhythms (Jaspers et al., 2015; Mortola, 2007; Yu et al., 2015). Although diverse proteins, such as HIF1A and mTOR, appear to interfere with the clock transcriptional regulation in low oxygen conditions (Walton et al., 2018; Wu et al., 2017), our data suggest that perturbation of MAX/MNT complexes might provide an essential contribution to hypoxia-induced chronodisruption.

In addition, genomic inactivation of MAX has been recently associated with a MYC-independent progression to malignancy of gastrointestinal stromal tumors (Schaefer et al., 2017) and future studies might reveal the contribution of the clock function of MAX in its paradoxical tumor suppressor role.

## Acknowledgments

This work was supported in part by the Associazione Italiana per la Ricerca sul Cancro (AIRC); grant number IG 17005. DNA and RNA sequencing was performed at the Genomic Unit of Center for Genomic Science, Fondazione Istituto Italiano di Tecnologia (IIT), Italy. We thank Dr Salvatore Bianchi for his valuable technical assistance with DNA and RNA library preparation, sequencing and quality control and Dr Alessio Ferrari for helping with generation of reporter cell lines. We also acknowledge the precious help with English corrections of Dr Colin J. Moore.

## Author Contributions

O.B., N.B., D.V., and A.G. performed and analyzed knockdown and qRT-PCR analyses. O.B. and D.V. performed and analyzed immunoblot analyses. O.B. performed and analyzed qChIP experiments and generated DNA and RNA libraries for sequencing. N.B. generated reporter cell lines, performed and analyzed real-time bioluminescence monitoring of circadian oscillation. O.C., S.C. and B.G. analyzed ChIP and RNA sequencing data. B.G. conceived the project and supervised experiments. O.B., S.C. and B.G. wrote the paper.

## Contact for Reagent and Resource Sharing

Further information and requests for resources and reagents should be directed to and will be fulfilled by the Lead Contact,Benedetto Grimaldi.

## Method Details

### Cell culture

Human breast cancer MDA-MB-231, BT549, human skin cancer A375, human stomach cancer SNU16, human embryonic kidney HEK-293 and HEK-293T cell lines (previously obtained from American Type Culture Collection (ATCC)) were grown in DMEM medium (Sigma, catalog#D8537) supplemented with 4 mM L-glutamine (EuroClone, catalog#ECB3000D), 10% fetal bovine serum (FBS) (Sigma, catalog#10001432) and 1X penicillin:streptomycin solution (Sigma, catalog#P4333).

Human liver cancer HEP-G2 cells (kindly provided by Istituto di Ricerche Farmacologiche ‘Mario Negri’, Milan, Italy) were grown in RPMI-1640 medium (EuroClone, catalog#ECB90006L) supplemented with 4 mM L-glutamine and 10% FBS.

Human epithelial MCF10A cell line (obtained from ATCC) were maintained in a 1:1 mixture of DMEM and Ham’s F12 (Sigma, catalog#51651C) media supplemented with 20 ng/ml human Epidermal Growth Factor (EGF), (Sigma, catalog#E9644), 2mM L-glutamine, 5% horse serum (Sigma, catalog#H1270), 10 μg/ml human recombinant insulin (Sigma, catalog#I9278), 0.5 mg/ml hydrocortisone (Sigma, catalog#H6909), 100 ng/ml cholera toxin (Sigma, catalog#C8052) and 1X penicillin/streptomycin solution.

Human foreskin fibroblast BJ-5ta cell line (ATCC, catalog#CRL-4001) were maintained in a 4:1 mixture of DMEM and 199 (Sigma, catalog#M3769) media supplemented with 10% FBS, 2mM L-glutamine and 1X penicillin/streptomycin solution.

All cell lines were maintained at 37◻°C in a humidified atmosphere with 5% CO_2_.

### siRNA transfection

For RNAi experiments, 30 nM siRNA sequences against *MAX*, *MYC*, *BMAL1*, *CRY1*, *CRY2*, and *MNT* were reverse-transfected with DharmaFect 1 Transfection reagent (Dharmacon, catalog#T-2001-03) following the manufacturer’s protocol. As a control, cells were transfected with MISSION® siRNA Universal Negative Control #1 (Sigma, catalog#SIC001).

### Cell proliferation analysis

MDA-MB-231 cells were transfected with siRNA sequence against *BMAL1*, *MAX*, *MYC* or a non-coding control. The number of cells was counted 24, 48, 72 and 96 hours after transfection with a Countess II FL (Life Technologies). Trypan blue staining was used to discriminate live and dead cells.

### Quantitative RT-PCR

RNA and cDNA samples were prepared by Trizol (Life Technologies, catalog#15596018) extraction and retro-transcription with SuperScript ViloTM Master Mix (Invitrogen, catalog#11755-250) following the manufacturer’s protocol. Relative transcript expression levels were assessed by quantitative PCR with iTaqTM Universal SYBR Green Supermix (BioRad, catalog#172-5124) on a Via7 thermocycler (Invitrogen). GAPDH transcripts were used for normalization. Primer sequences are listed in Supplemental Table 6.

### RNA sequencing

Total RNA samples were prepared by Trizol extraction followed by purification with PureLink RNA kit (Invitrogen, catalog#12183018A). RNA integrity was examined using capillary electrophoresis on a BioAnalyzer 2100 (Agilent Technologies, USA). RNA libraries for sequencing were prepared with Illumina RNA TruSeq kit v2 (Illumina, catalog#15027084) following the manufacturer’s protocol. Libraries were sequenced using 50 base pairs paired-end on an Illumina HiSeq 2000 sequencer. RNA-Seq reads were aligned with tophat v.2.0.8 with -r 170 -p 8 --no-novel-juncs --no-novel-indels --librarytype fr-unstranded options (Kim et al., 2013).

### Analysis of differentially expressed genes

RNA-seq counts were used to determine differentially expressed genes with DeSeq2 package (https://doi.org/10.1186/s13059-014-0550-8) included in the Galaxy web platform (usergalaxy.org). Differentially expressed genes (DEGs) were defined adopting an adjust P < 0.001 as a statistical cut-off value. Lists of DEGs in BMAL1 and MAX-silenced cells, logarithmic fold change (LogFC), adjust P values (adjP) and normalized counts are provided in Supplemental Table 3-4. Analysis for co-occurrence in common KEGG pathways was Molecular Signatures Database v6.2 package using a False Discovery Rate (FDR) < 0.01. Output normalized counts from siBMAL1, siMAX and control samples were used to generate heat map and clustering analysis of common DEGs in BMAL1 and MAX-silenced cells with Morpheus online software (https://software.broadinstitute.org/morpheus/). Only genes with an absolute logarithmic fold change > 0.5 were included in this analysis.

### Quantitative chromatin immunoprecipitation

Chromatin immunoprecipitation (ChIP) experiments in MDA-MB-231-silenced cells were performed 48 hours after reverse-transfection with siRNA sequences. ChIP experiments in MCF10A were performed on 80% confluent cells. Cells were crosslinked for 10 min with 1% formaldehyde (Sigma, catalog#F8775), neutralized with 125 mM glycine at pH 2.5 for 5 min and washed twice in PBS. Cells were lysed with 0.5% SDS buffer containing protease inhibitors cocktail (SIGMA, catalog# P8340), scraped and centrifuged at 1150 g for 10 min at 4◻C. Pellets were resuspended in 4 ml of ice-cold IP Buffer composed by a 2:1 micture of SDS buffer (100 mM NaCl, 50mM Tris-Cl, pH8.1 EDTA, pH 8.0, 0.5% SDS) and Triton Dilution Buffer (100 mM Tris-Cl, pH 8.6, 100 mM NaCl, 5mM EDTA, pH 8.0, 5% Triton X-100) containing protease inhibitor cocktail. Samples were sonicated with Branson Digital Sonifier (Danbury, USA) in 30s bursts followed by 30s of cooling on ice for a total sonication time of six minutes per sample. Chromatin was pre-cleared for 1 h at 4 ◻C with Sepharose protein G beads (Life technologies, catalog# 101242) and subsequently precipitated overnight at 4 ◻C with 4 μg of anti-BMAL1 (Protein Tech, catalog# 14268-1-AP), 4 μg of anti-MAX (Bethyl Lab, catalog#A302-866A), 4 μg of anti-MNT (Bethyl Lab, catalog#A303-626A) 4 μg of anti-MYC (Cell Signaling catalog#13987S) and 4 μg of normal Rabbit IgG (Cell Signaling #2729S) as a negative control. DNA protein complexes were recovered with Sepharose protein G beads overnight and washed twice sequentially with Mixed Micelle Wash Buffer (150 mM NaCl, 20 mM Tris-Cl, pH 8, 5 mM EDTA, 5% w/v sucrose, 1% Triton X-100, 0.2% SDS), LiCl/Detergent buffer (0.5% Na-deoxycholate, 1 mM EDTA, 250 mM LiCl, 0.5% (v/v) NP-40, 10 mM Tris-Cl, pH 8.0), Buffer 500 (0.1% (w/v) deoxycholic acid, 1 mM EDTA, 50 mM HEPES, pH 7.5, 500 mM NaCl, 1% (v/v) Triton X-100) and TE buffer (10mM Tris-Cl, 1mM EDTA, pH 8.0). Beads were further supended in TE-S buffer (TE buffer, 2% SDS) and treated with RNAse A for 30 min at 37 ◻C. Cross-linking was reverted by overnight incubation at 65 ◻C in TE-S containing 0.4 mg/ml proteinase K. In parallel, inputs were treated in the same way. Immunoprecipitated and input DNA was purified using PCR purification kit (QIAGEN, catalog#28106) using 60 μl of buffer T (10mM Tris-Cl, pH 8.0). Quantitative PCR was performed by using iTaqTM Universal SYBR Green Supermix. Primer sequences are listed in Supplemental Table S6. Promoter occupancy was calculated as percent of input using the following formula: [2^−(CT_ChIP_−CT_input_)] × [Input dilution factor].

### Chromatin immunoprecipitation sequencing

For ChIP sequencing, immunoprecipitated DNA from MDA-MB-231 chromatin samples was obtained with the protocol described for quantitavie ChIP. Input and immunoprecipitated DNA (1–10 ng) were blunt-ended and phosphorylated, and a single ‘A’ nucleotide was added to the 3’ ends of the fragments in preparation for ligation to an adapter that has a single-base ‘T’ overhang. The ligation products was purified and accurately size-selected by agencourt AMPure XP beads (Beckman Coulter, catalog# A63881). Purified DNA was PCR-amplified to enrich for fragments that have adapters on both ends. All the steps were performed on automation instrument, Biomek FX by Beckman Coulter. The final purified product was then quantitated prior to cluster generation on bioanalyzer 2100. Libraries with distinct adapter indexes were multiplexed (1/5 libraries per lane) and after cluster generation on FlowCell were sequenced for 50 bases in the single read mode on a HiSeq 2000 sequencer.

### ChIPseq data analysis

Alignments of reads and peak calling were performed using HTS-flow (Bianchi et al., 2016). Brefly, ChIP-Seq reads were aligned on hg19 human genome assembly using BWA v.0.6.2 (Li and Durbin, 2009) and peaks were called with MACS2 v.2.0.9 (Feng et al., 2012), using a p-value threshold of 10^-8. The normalized reads count in genomic regions (rpm) and plots were obtained using custom R scripts (Team, 2013). Peaks of ChIP-Seq were considered to belong to a promoter if at least 1 bp of the peak was contained in the [−300;+300] interval from TSS (transcription start sites). Promoters were retrieved from UCSC using TxDb.Mmusculus.UCSC.hg19.knownGene R package (Huber et al., 2015).

### Immunoblotting

Protein samples were extracted in RIPA buffer as described previously (De Mei et al., 2015). Immunoblot were performed on 20◻μg of protein extracts separated on 8-15% polyacrylamide gels and transferred onto nitrocellulose membrane (GeHealthcare Life Science, catalog# 10600001). Immunoblot signals were visualized by the chemiluminescent ECL Star substrate (Euroclone, catalog# EMP001005) using the ImageQuant LAS-4000 Chemiluminescence and Fluorescence Imaging System (Fujitsu Life Science, Japan). Densitometry analysis was performed with ImageJ software (Wayne Rasband, USA).

### Generation of PER2 promoter reporter plasmids

A synthetic 778 base-pair DNA fragment corresponding to the wild-type human *PER2* promoter (from 238288785 to 238289562 of *Homo sapiens* chromosome 2, GRCh38.p12 primary assembly, sequence ID: NC_000002.12) and a corresponding fragment containing the E’-box sequence CACGTT mutated in CCCCCC were synthetized by and cloned in pBluescript II KS (−) vector by GenScript (USA). Wild-type and mutated PER2 promoter sequences were then sub-cloned in pGreenFire1-mCMV (EF1α-neo) (System Biosciences, catalog# TR010PA-N) upstream the minimal CMV promoter driving the expression of the copGFP protein. The resulting E’-box-GFP and E’^mut^-box-GFP vectors were sequenced with a specific primers in reverse orientation respect with the *copGFP* gene (copGFP RV primer, 5’-GATGATCTTGTCGGTGAAGATCACG-3’) to confirm the correct cloning.

### Generation of reporter cell lines

BMAL1:luc (obtained from Addgene, plasmid# 46824), E’-box-GFP and E’mut-box-GFP lentiviral vectors were packed in HEK293T cells by co-transfection with pVSV-G and pSAX2 plasmids (obtained by addgene, plasmid# 8454 and plasmid# 12260). Supernatant were collected 48 hours after transfection and concentrated by ultracentrifugation. For lentiviral infection, MDA-MB-231 and MCF10A cells were exposed to lentiviral particle for 48 h prior to be moved to a selective medium containing either 1 μg/ml of puromycin (Sigma-Aldrich, catalog# P8833, BMAL1:luc) or 600ug/ml of G418 (Euroclone S.p.A, catalog# ECM0015Z, E’-box-GFP and E’^mut^-box-GFP). Single cell clones were selected via limiting dilution, and the expression of luciferase and GFP reporters was confirmed by luminescence and fluorescent microscopy analyses, respectively.

### Real-time bioluminescence monitoring of circadian rhythm in cultured cells

MDA-MB-231 and MFC10A cells expressing the circadian reporter BMAL1:luc were transfected with siRNA against BMAL1, MAX, MNT or a non-coding control. Fourty-eight hours post transfection, cells were synchronized by a treatment with 500 nM dexamethasone (dex) for 2 h. After the replacement of dex-containing mediuam with a warm phenol red-free medium supplemented with 0.4 mM D-luciferin (Invitrogen, catalog#L2916), cells were placed into a real-time bioluminescence reader (LumiCycle36, Actimetrics, Wilmette, IL, USA) maintained at 37◻°C in a humidified atmosphere with 5% CO_2_. Luminescence was recoreded every 5 minutes over a 4-6 days period. To compare rhythmic patterns in-silenced cells, baseline-subtracted luminescence data were fitted to a sine wave using LumiCycle analysis software (Actimetrics, Wilmette, IL, USA). For circadian analysis in MCF10A, a medium free of horse serum and hydrocortisone was used during synchronization by dexamethasone treatment and luminescence monitoring.

### Sequencing Data availability

The datasets and computer code produced in this study are available at Gene Expression Omnibus (https://www.ncbi.nlm.nih.gov/geo/query/acc.cgi?acc=GSE127192), accession numbers GSE127192 and GSE127212.

### Statistical Analysis

For qRT-PCR, qChIP and cell proliferation analyses, statistical significance between groups was calculated by two-way ANOVA associated with Bonferroni post-tests. The correlation between BMAL1/MAX-bound DEGs genes was evluated using Spearman’s rank correlation test. These statistical analyses were performed using Prism6 software package. Significance values were P < 0.05 (*), P < 0.01 (**) and P < 0.001 (***). For the analysis of of DEGs, Benjamini-Hochberg adjusted p value (adjP) was calculated using DeSeq2 package and was considered significant at adjP < 0.05. Co-occurrence analysis of DEGs in KEGG pathways was performed with Molecular Signatures Database v6.2 package (Broad Institute, http://software.broadinstitute.org/gsea/msigdb/index.jspof) using a False Discovery Rate (FDR) q value < 0.01 as a statistical cut-off.

