## Supplementary figures and images for "MYC-associated factor MAX is an essential regulator of the clock core network"

### Figure S1

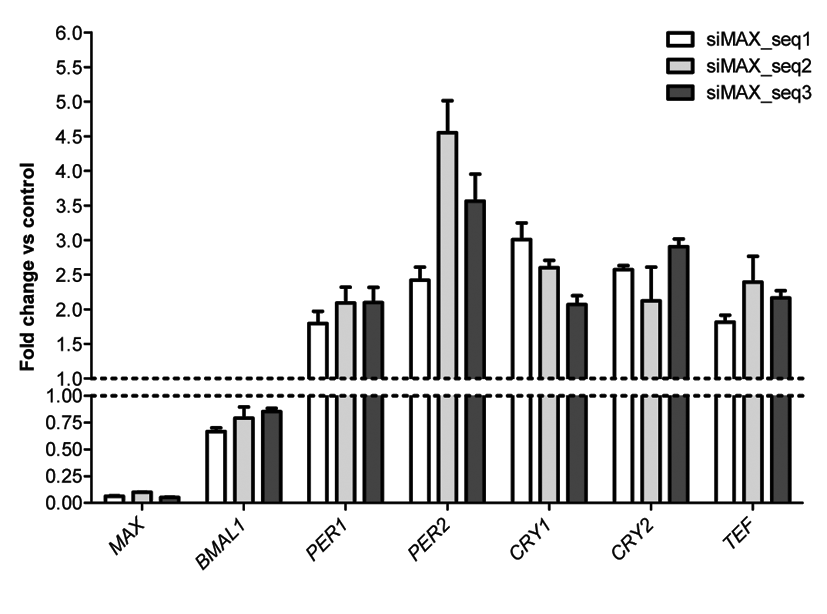

### Figure S2

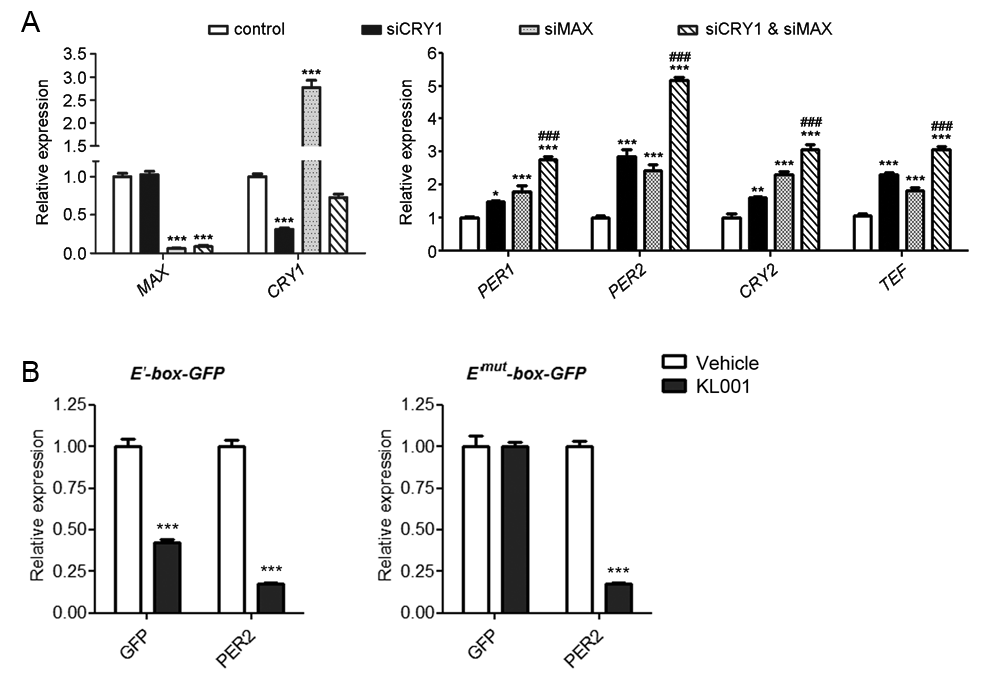

### Figure S3

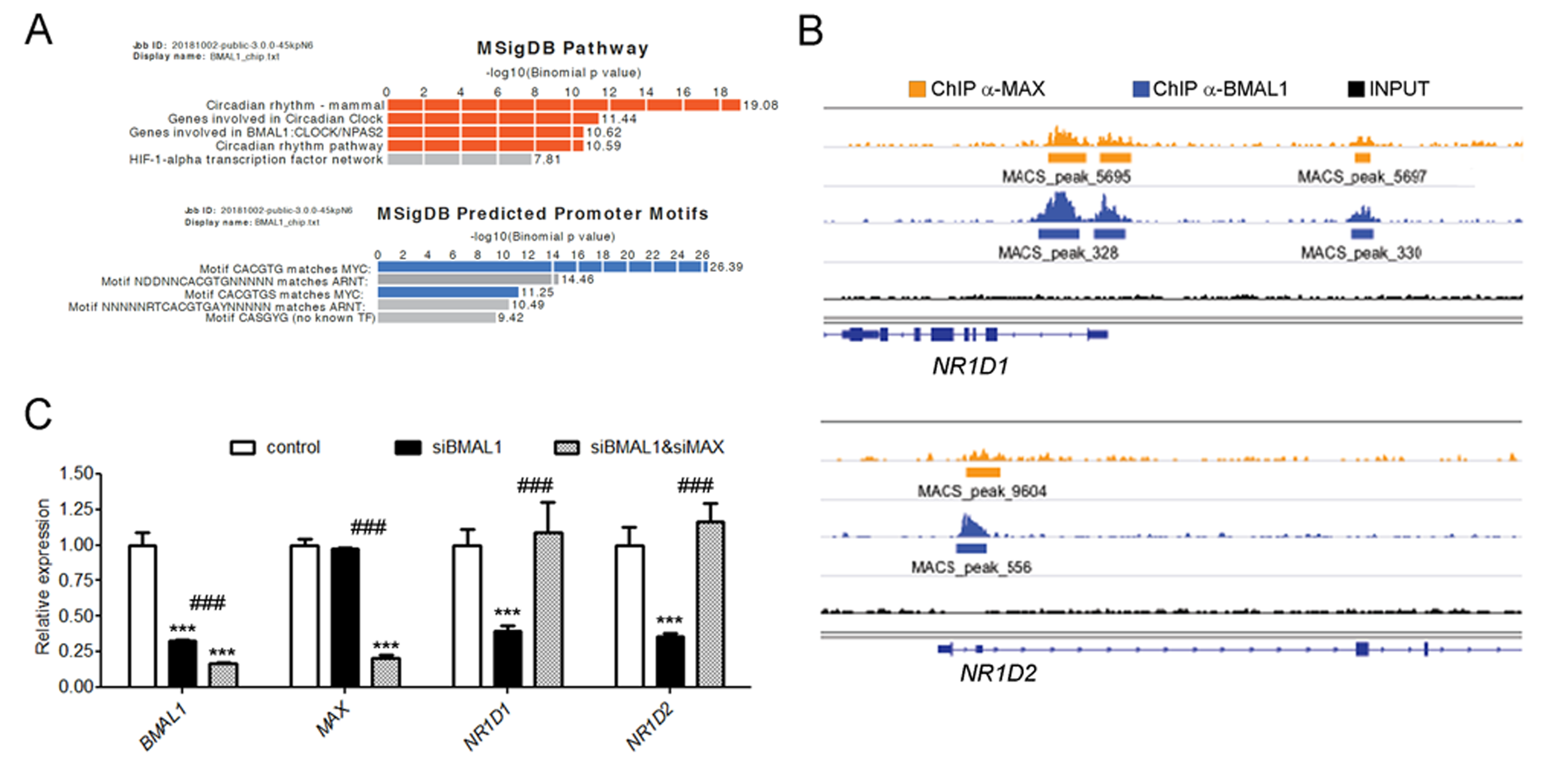

### Figure S4

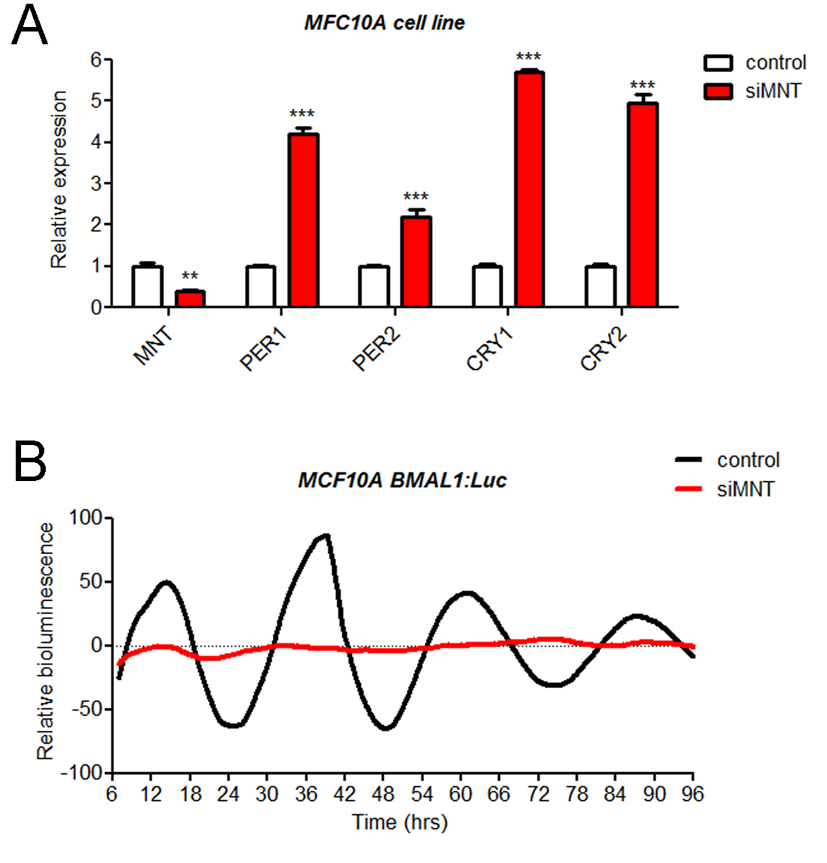
