## supplemental info for "MYC-associated factor MAX is an essential regulator of the clock core network"

**Supplementary information**

**Supplementary Figure legends**

**Figure S1, related to Figure 1.** Specificity of MAX silencing on the expression of clock core genes. Expression of the indicated clock genes in MDA-MB-231 cells upon knockdown of MAX with three diverse and non-redundant siRNA sequences. A non-coding siRNA was used as control. Relative expression was determined by qRT-PCR using *GAPDH* for normalization. Values of control cells were set to 1. Shown as mean fold change versus control + SEM, n = 3.

**Figure S2, related to Figure 2.** (*A*) Expression of clock genes (*PER1*, *PER2*, *CRY2*, and *TEF*) in MDA-MB-231 cells with knocked down *CRY1* (siCRY1), *MAX* (siMAX), and both *CRY1* and *MAX* (siCRY1&siMAX). A non-coding siRNA was used as control. Relative expression was determined by qRT-PCR using *GAPDH* for normalization. Values of control cells were set to 1. Shown as mean ± SEM, n ≥ 3. *P < 0.05. **P < 0.01 and ***P < 0.001, two-way ANOVA with Bonferroni post hoc test, silencing versus control. ^###^P < 0.001, siCRY1&siMAX versus siCRY1. (*B)* Repression of *PER2*-promoter driven *GFP* by a CRY agonist, KL001 (5 µM), in reporter MDA-MB-231 cell lines bearing either a wild type (*E’-box-GFP*) or a mutated (*E^’mut^-box-GFP*) PER2 promoter sequence. Expression of the endogenous *PER2* was used as a control of the responsiveness of the cells to KL001. Two-way ANOVA with Bonferroni post hoc test comparing KL001 versus vehicle. **P < 0.01 and ***P < 0.001.

**Figure S3, related to Figure 3.** (*A*) Ontological annotation of BMAL1 bound regions by Genomic Regions Enrichment of Annotations Tool (GREAT) (http://great.stanford.edu/public/html/). A significant enrichment for circadian regulated genes and E-box motifs is shown. (*B*) Genomic snapshots of the promoter region of the circadian repressor genes, *NR1D1* (also known as *REV-ERBα*) and *NR1D2* (also known as *REV-ERBβ*) showing the enrichment of MAX (orange) and BMAL1 (blue) ChIP-seq signals. (*C*) Expression of *NR1D1* and *NR1D2* in MDA-MB-231 cells upon the knockdown of either BMAL1 (siBMAL1) or BMAL1 and MAX (siBMAL1&siMAX). Relative expression was determined by qRT-PCR using *GAPDH* for normalization. Values of control cells were set to 1. Shown as mean fold change versus control + SEM, n = 3. Two-way ANOVA with Bonferroni post hoc test is shown. ***P < 0.001, silenced versus control cells, and ^###^P < 0.001, MAX-silenced versus BMAL1-silenced cells.

**Figure S4, related to Figure 7.** MNT controls circadian gene expression in MCF10A cells. (*A*) Expression of *MNT*, *PER1*, *PER2*, *CRY1*, and *CRY2* genes upon MNT silencing in epithelial MCF10A cells. Relative expression was determined by qRT-PCR using *GAPDH* for normalization. Values of control cells were set to 1. Shown as mean + SEM, n ≥ 3. Two-way ANOVA with Bonferroni post hoc test is shown. **P < 0.01 and ***P < 0.001, silenced versus control cells. (*B*) Real-time bioluminescence oscillatory pattern in MNT-silenced and control MCF10A cells expressing a firefly luciferase reporter controlled by the circadian-responsive *BMAL1* promoter. Shown as baseline-subtracted luminescence data fitted to a sine wave.

**Supplementary Tables**

**Table S1**

Genomic location of ChIP-seq peaks from MDA-MB-231 chromatin samples immunoprecipitated with α-MAX antibody.

**Table S2**

Genomic location of ChIP-seq peaks from MDA-MB-231 chromatin samples immunoprecipitated with α-BMAL1 antibody.

**Table S3**

Differentially Expressed Genes (DEGs) in BMAL1 silenced MDA-MB-231 cells compared with control. Mean normalized counts (Mean), log fold change difference (LogFC) and adjusted P values (adjP) are reported.

**Table S4**

Differentially Expressed Genes (DEGs) in MAX silenced MDA-MB-231 cells compared with control. Mean normalized counts (Mean), log fold change difference (LogFC) and adjusted P values (adjP) are reported. Related to Fig. 4.

**Table S5**

Genes significantly affected by the knockdown of either MAX or BMAL1 (adjust P < 0.05). Log fold change difference (LogFC) in MAX-silenced and BMAL1-silenced cells compared with control is reporter. Normalized counts of triplicates from siMAX, siBMAL1 and control samples is also reported. These values were used for generating the heat map in Fig. 4C.
